# PARNAS: Objectively Selecting the Most Representative Taxa on a Phylogeny

**DOI:** 10.1101/2022.09.12.507613

**Authors:** Alexey Markin, Sanket Wagle, Siddhant Grover, Amy L. Vincent Baker, Oliver Eulenstein, Tavis K. Anderson

## Abstract

The use of next-generation sequencing technology has enabled phylogenetic studies with hundreds of thousands of taxa. Such large-scale phylogenies have become a critical component in genomic epidemiology in pathogens such as SARS-CoV-2 and influenza A virus. However, detailed phenotypic characterization of pathogens or generating a computationally tractable dataset for detailed phylogenetic analyses requires bias free subsampling of taxa. To address this need, we propose *parnas*, an objective and flexible algorithm to sample and select taxa that best represent observed diversity by solving a generalized k-medoids problem on a phylogenetic tree. *parnas* solves this problem efficiently and exactly by novel optimizations and adapting algorithms from operations research. For more nuanced selections, taxa can be weighted with metadata or genetic sequence parameters, and the pool of potential representatives can be user-constrained. Motivated by influenza A virus genomic surveillance and vaccine design, *parnas* can be applied to identify representative taxa that optimally cover the diversity in a phylogeny within a specified distance radius. We demonstrated that *parnas* is more efficient and flexible than current approaches, and applied it to select representative influenza A virus in swine genes derived from over 5 years of genomic surveillance data. Our objective selection of 4 to 6 strains selected every two years from the 16 distinct genetic clades were sufficient to cover 80% of diversity circulating in US swine. We suggest that this method, through the objective selection of representatives in a phylogeny, provides criteria for rational multivalent vaccine design and for quantifying diversity. PARNAS is available at https://github.com/flu-crew/parnas.

## 1 Introduction

Next-generation sequencing technologies are routinely applied to generate thousands to millions of genomes. From the beginning of the COVID-19 pandemic to present, more than 11 million SARS-CoV-2 viruses have been sequenced (Turakhia *et al*., 2021). Genomic epidemiology and phylogenetic analysis are essential tools to navigate this landscape of genetic data (Hill *et al*., 2021). However, phylogenetic trees are often insufficient to make informed intervention decisions, especially in public health. For example, it is difficult to select a few representative virus strains from within a phylogeny to include in a polyvalent vaccine; and selecting unique pathogen strains to represent variable genes or genomes for detailed *in vivo* studies on transmission and pathology requires objective criteria that are reproducible. As these assays cannot be performed on the same scale as sequencing, bias-free subsampling strategies are necessary. Subsampling techniques must satisfy three properties: (i) selected taxa should capture observed genetic diversity; (ii) selected taxa should be representative of respective diversity groups; and (iii) the method needs to be flexible to allow preferential weighting of taxa and be open to specific constraints, such as a desire to target spatial or temporal metadata due to limited availability of some strains for characterization.

We consider the above problem as representative sampling and formulate it as the *k*-medoids problem on a phylogenetic tree. That is, the goal is to select *k* representative taxa, such that the overall distance from all taxa to the respective closest representative is minimized. This formulation simultaneously partitions the taxa into *k* (phylo)genetic clusters and chooses best representatives within the clusters, thus satisfying properties (i) and (ii) from above. This problem was originally considered in the context of biodiversity (Faith, 1994) and later as a means to improve phylogenetic inference (*Matsen et al*., 2013). We show that this problem is efficiently solvable by adapting an algorithm for a related problem in operations research (Tamir, 1996). We implemented this algorithm with additional computational optimizations in the software package *parnas* (https://github.com/flu-crew/parnas). We also developed *parnas* to solve a more general problem and satisfy property (iii) by allowing taxa to be weighted by metadata so that those with larger weights are better represented within the selection process. Additionally, in *parnas*, users can constrain the pool of potential representatives in multiple ways and indicate previously selected/employed representative taxa, so that new representatives will optimally complement prior selections.

In virology, the diversity of viruses continually changes through evolutionary processes such as mutation and selection, and it is necessary to monitor and identify possible emerging threats. *parnas* can be applied to identify genetically unique pathogens by allowing users to specify a ‘coverage radius,’ such that each potential representative covers all diversity within the specified radius (evolutionary distance) on the phylogenetic tree. Thus, it is possible to choose *k* representatives so that the total amount of covered diversity is maximized, or choose the minimum number of representatives that cover all the diversity on a tree. The coverage problem also allows the optimal elimination of redundancy from a tree if a small radius is specified. Though linking genetic diversity to phenotype is challenging (Zeller *et al*., 2021), distance thresholds have been applied to identify discrete genetic clades (Han *et al*., 2019) that are correlated with antigenic differences (Anderson *et al*., 2020). Therefore, finding optimal genetic coverage on a phylogenetic tree can identify groups of viruses that may be phenotypically novel and drifted from existing viruses.

In this study, we developed an algorithm that solves the *k*-medoids problem on a phylogenetic tree. We compared the runtime of *parnas* to prior approaches for the (unweighted) *k*-medoids problem on a phylogenetic tree proposed by Matsen *et al*., 2013 as the ADCL algorithms (ADCL-PAM and ADCL-full). Despite ADCL-PAM being an inexact heuristic, we observed *parnas* to be more scalable than both ADCL-PAM and the exact ADCL-full algorithm. We demonstrated that novel optimizations introduced in *parnas* reduce the dynamic programming table size and resulted in 99.6% performance improvements in terms of runtime and RAM. Further, we demonstrated that *parnas* is faster than Treemmer (Menardo *et al*., 2018), a popular tool for eliminating redundancy on a phylogeny, while preserving diversity. We showed that Treemmer taxon selections can be 40-50% less representative than the optimal selections by *parnas*.

Then we applied *parnas* to an empirical influenza A virus (IAV) in swine dataset derived from the national USDA IAV in swine surveillance system (Anderson *et al*., 2013). Our goal was to determine a minimum number of representatives that maximized genetic coverage, and assess the duration that the representatives covered a significant amount of diversity over time.

## 2 Materials and Methods

### 2.1 Definitions

A *(phylogenetic) tree* over a *taxon set L* is a rooted and full binary tree *T* = (*V, E*) with its leaves uniquely labeled (i.e., identified) by the elements of *L* and *edge lengths* described by the function *l* : *E* → ℝ^+^. The root of *T* is denoted by *ρ*(*T*). For a node *v* in *T* we denote its parent by *p*(*v*) and its children by *v*_(1)_ and *v*_(2)_ if such nodes exist. By *T*_*v*_ we denote the subtree of *T* rooted at *v*.

The *distance* between two nodes *u, v* in *T*, denoted by *d*(*u, v*), is defined as the sum of the edge lengths along the simple path between *u* and *v*.

To allow for unrooted and/or multifurcating trees, given such a tree, we arbitrarily root it and add the minimum number of edges with length 0 to make it strictly bifurcating and preserve pairwise taxon distances.

### 2.2 Representative sampling

Our core problem is as follows: given a tree *T*, identify a set of its leaves that best represents its taxon diversity. Problem 1 formalizes this idea. We allow leaves in *T* to be weighted according to some real-valued function *w* : *L* → ℝ^+^ (the default weights are *w*(*l*) = 1 for all *l ∈ L*).

#### Problem 1.

*Given a tree T over a taxon set L, and a positive integer k <* |*L*|, *find*

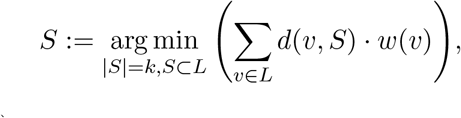

*where d*(*v, S*) := min_*c∈S*_ *d*(*v, c*).

That is, a set of representatives *S* should minimize the sum of weighted distances from all leaves to their closest representatives. This problem is a weighted version on the famous k-medoid problem (Kaufman and Rousseeuw, 1990) on a phylogenetic tree.

Next, we want to account for potentially pre-selected representatives *C* and a coverage radius *r*. The coverage radius implies that a single representative covers all the diversity on the tree within the radius. Therefore, we define the function

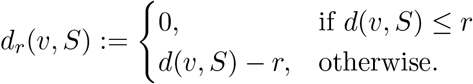

Problem 2 then generalizes Problem 1 and accounts for *C* and *r*.

#### Problem 2.

*Given a positive integer k <* |*L*|, *a non-negative radius r, and a set C ⊂ L of prior representatives, find*

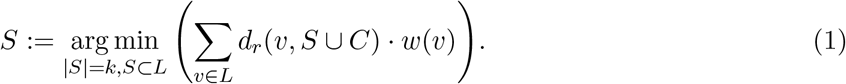

We also define *d*_*r,C,w*_(*v, S*) := *d*_*r*_(*v, S ∪ C*)*w*(*v*) to simplify notation.

**Proposition 1**. *Problem 2 can be solved in O*(*n*^2^*k*) *time for n* = |*L*| *using Tamir’s algorithm for the p-medians problem on a tree*.

*Proof*. Tamir, 1996 solved the following generalization of the p-medians problem on trees:

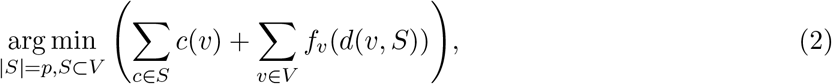

where *c*(*v*) is some cost function on the nodes of the tree and *f*_*v*_ is a non-negative and non-decreasing function specific to node *v*.

We claim that Problem 2 is a special case of generalized p-medians. We show the reduction by first assigning

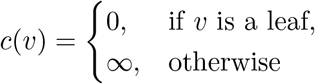

to prevent selecting non-leaf nodes as representatives. Further, we set *f*_*v*_ to 0 for all non-leaf *v*. For *v ∈ L*, let *m*_*v*_ := *d*(*v, C*) if *C ∅* and *m*_*v*_ := *∞* otherwise. Then

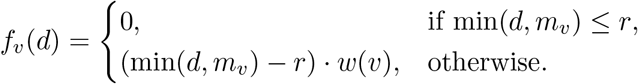

Observe that for *v ∈ L, d*_*r*_(*v, S ∪ C*) *· w*(*v*) = *f*_*v*_(*d*(*v, S*)). It is then apparent that under this reduction Eq. (2) is equivalent to Eq. (1) when *S ⊂ L* and *p* = *k* (recall that *S ⊂ L* is enforced by cost assignments).

### 2.3 Improving the dynamic programming solution for Problem 2

Tamir’s dynamic programming algorithm for generalized p-medians has both best-case and worst-case time and space complexity of *O*(*kn*^2^). This can be limiting in practice for trees with over 1,000 leaves and large *k*. Therefore, we show how to achieve better (in-practice) time and space efficiency.

Similar to Tamir, 1996, for *v* ∈ *T, q* ≤ *k*, and *r* ∈ *T*_*v*_ ∩ *L* we define *G*(*v, q, r*) as an optimal value of the subproblem of Problem 2 constrained to *T*_*v*_, *q* representatives, and a condition that there exists a representative that is at most *d*(*v, r*) away from *v*. Similarly, for *r* ∈ *T* \ *T*_*v*_, *F* (*v, q, r*) is an optimal value of the subproblem of Problem 2 constrained to *T*_*v*_, *q* representatives, and with a knowledge that there exists a representative in *T* \ *T*_*v*_ on distance *d*(*v, r*) away from *v*. Observation 1 and Lemma 2 remove redundancies in the dynamic programming.

#### Observation 1.

*Let s* = |*T*_*v*_ *∩ L*| *be the number of leaves in T*_*v*_. *Then for q > s, G*(*v, q, r*) = *G*(*v, s, r*) *and F* (*v, q, r*) = *F* (*v, s, r*), *since s is the maximum possible number of representatives in T*_*v*_.

#### Lemma 2.

*For q* > 0, *let r*^−*q*^ *denote the q-th farthest from v leaf in T*_*v*_, *and d*^−*q*^ := *d*(*v, r*^*−q*^). *Then for any r ∈ L with d*(*v, r*) > *d*^−*q*^,

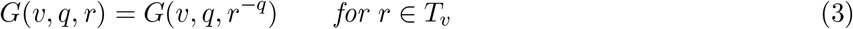

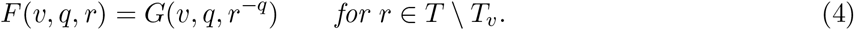

*Proof*. As there are at least *q* representatives in *T*_*v*_, at least one of them has to be no further then *d*^*−q*^ from *v*. Eq. (3) then follows from the definition of *G*.

Further, note that for *r ∈ T \ T*_*v*_ with *d*(*v, r*) *> d*^*−q*^ and any leaf *l* ∈ *T*_*v*_, we have *d*(*l, r*) = *d*(*l, v*)+*d*(*v, r*). Let *d*_*l*_ be the distance from *l* to the closest representative in *T*_*v*_. Then *d*_*l*_≤ *d*(*l, v*)+*d*^*−q*^. That is, any *l* is closer to a representative in *T*_*v*_ than to *r*. Hence, having *r* as a representative does not affect the subproblem in *T*_*v*_.

Lemma 2 implies that we do not need to compute subproblems *G*(*v, q, r*) and *F* (*v, q, r*) when *d*(*v, r*) > *d*^−*q*^. The above observations significantly improve the best-case complexity of the algorithm:

#### Lemma 3.

*The best-case time and space complexity of the dynamic programming algorithm is O*(min*{kn* log *n, n*^2^*}*), *which is achieved when T is a perfectly balanced tree with uniform edge lengths*.

In the Results section, we demonstrate that the above observations proved highly effective on simulated data.

### 2.4 Representative coverage

In addition to the representative sampling, we solve a coverage problem. Similarly to Problem 2, we are given (an optional) set of prior representatives *C* and a coverage radius *r*. The goal is to find a minimum set of representatives *S*, so that *S* ∪ *C* covers all leaves in the tree (within radius *r*).

#### Problem 3.

*Given set C ⊂ L and non-negative r ∈ ℝ, find minimum set S, s*.*t*.

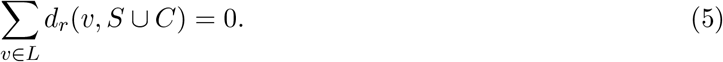

We show that this problem can be solved in *O*(*n*^2^) time using dynamic programming similar to Tamir, 1996.

For each node *v*_*i*_ with 1 *≤ i ≤* |*V* |, let 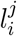 be the *j*-th farthest from *v*_*i*_ leaf in *L*. Ties are resolved, so that leaves in 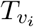 precede leaves outside the subtree; if two tied leaves are both within 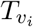 or both outside, then they are placed in order of the post-order traversal of *T*. The 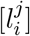 lists for each *v*_*i*_ can be computed in *O*(*n*^2^) time (Tamir, 1996).

We define 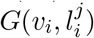 to be the minimum number of representatives required to cover 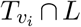, so that at least one of these representatives is 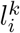, where *k ≤ j*. For 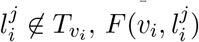 is the minimum number of representatives required to cover 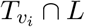 *∩ L*, while the closest representative outside of 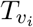 is 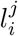.

**Base case**. If *v*_*i*_ is a leaf, 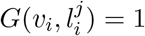 for all *j*. If *d*(*v*_*i*_, *C*) *≤ r* (i.e., *v*_*i*_ is covered by one of the prior centers), then 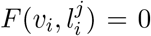 for all *j*. Otherwise, 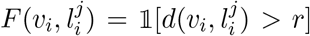 for all *j*, where 𝟙 is the indicator function.

**Internal nodes**. For non-leaf *v*_*i*_ and 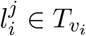, let *v*_*i*(1)_ and *v*_*i*(2)_ denote the children of *v*_*i*_. For 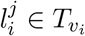, WLOG assume that 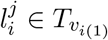, then

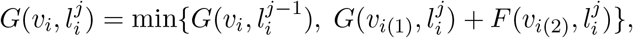

Where 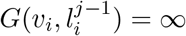 for *j* = 0. Further, for 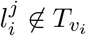, we have

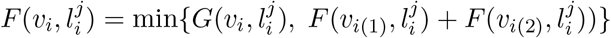

The optimal number of representatives required to cover *L* is then given by *G*(*ρ*(*T*), 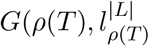. Consequently, the algorithm runs in *O*(*n*^2^) time and space.

### 2.5 Constraining the pool of representatives

We allow users to add constraints of two types to Problems 2 and 3:

i. A subset of taxa *A* can be chosen as representatives, but do not contribute to the objective function (i.e., excluded from summation in Equations 1 and 5).
ii. A subset of taxa *B* contribute to the objective function, but cannot be chosen as representatives.

For Problem 2, both constraints can be satisfied by appropriately assigning functions *c* and *f*_*v*_ for the respective taxa. In particular, (i) is satisfied by assigning *f*_*v*_ = 0 for *v ∈ A* and (ii) is satisfied by assigning *c*(*v*) = *∞* for *v ∈ B*.

Problem 3 with the added constraints can be solved in a similar fashion. However, it is possible that inclusion of constraint (ii) will make the complete coverage infeasible. In that case, *parnas* finds representatives that cover as much diversity as possible by iteratively solving Problem 2 with increasing the number of representatives *k*, until the objective function cannot be further improved.

### 2.6 Variations of the coverage radius

When using a maximum likelihood phylogenetic tree topology constructed from nucleotide sequences, one can be interested in re-scaling the tree so that branch lengths represent the number of amino acid substitutions. This can be needed to appropriately specify a coverage radius in relation to % amino acid sequence divergence rather than nucleotide divergence. *parnas* offers this option by using a user-specified amino acid alignment and TreeTime (Sagulenko *et al*., 2018) to infer ancestral amino acid substitutions and rescale the tree edges to represent the number of substitutions on that edge. This is motivated by the influenza A virus analysis, where specifying a 5% amino acid divergence radius for HA1 subunit has direct applications in vaccine strain selection and identification of antigenically novel viruses (Anderson *et al*., 2020).

Additionally, we implemented a *binary* coverage problem, where *parnas* selects representatives that cover as much taxa weight as possible (instead of weighted diversity), as follows:

#### Problem 4.

*Given a positive integer k <* |*L*|, *a non-negative radius r, and a set C ⊂ L of prior representatives, find*

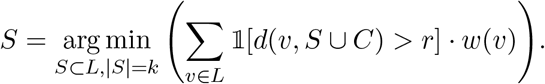

### 2.7 Quantifying represented diversity

Given a solution to Problem 2, we may ask how much diversity the selected strains represent. To help answer this question, we adapt ideas from k-means clustering, where an optimal number of clusters is often chosen based on the ‘variance explained’ by the cluster centers – a concept related to an F-ratio statistic (Bock, 1985). In particular, given a set *X ∈* ∝^*p*^ and a partition of *X* into clusters *X*_1_,…, *X*_k_ with cluster means 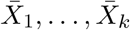, the variance explained can be calculated as

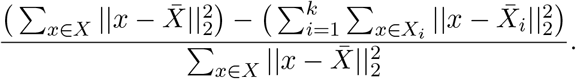

That is, the total sum of squares (TSS) minus within-cluster sum of squares (WSS), divided by TSS. Our computational problem is defined in terms of representatives. Hence, we define ‘cluster means’ as their respective representatives. That is, let *m*_0_ be a representative for entire set *L* (a solution to Problem 2 for *k* = 1). Further, let *S* = *{m*_1_, …, *m*_*k*_*}* be a solution to Problem 2 for *k >* 1. We partition set *L* into *L*_1_, …, *L*_*k*_ according to the closest representatives *m*_*i*_. Then, we say that *S represents P* % *of overall diversity*, where

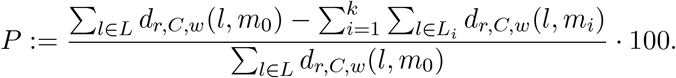

Note that *P* accounts for the coverage radius *r* and prior representatives *C*. Intuitively, each selected leaf represents the leaves closest to it. Then *d*_*r,C,w*_(*l, m*) is an ‘error’-term for leaf *l. P* measures the reduction in error from 1 to *k* representatives and hence the increase in represented (weighted) diversity.

### 2.8 Runtime comparison with ADCL

We implemented *parnas* using Python 3 and Numba (Lam *et al*., 2015) for just-in-time compilation of the dynamic programming algorithms. We simulated 80 birth-death trees with the birth rate *µ* = 1 and death rate *δ* = 0.5 and the number of leaves, *n*, varying between 500 and 4000 with step 500 (10 trees per fixed *n*). We then executed *parnas*, ADCL-full, and ADCL-PAM on each tree with fixed *k* = 100 (i.e., choosing 100 representatives). ADCL-full and ADCL-PAM are methods from (Matsen *et al*., 2013) which address Problem 1 and are a part of the *pplacer* suit of methods (Matsen *et al*., 2010). We measured the runtime of each method as well as the memory savings achieved in *parnas*.

Additionally, we evaluated how the methods scale as the number of representatives (*k*) grows. We ran all three methods on the trees with *n* = 2000 leaves and *k* varying from 40 to 1000 with step 40. This study was conducted on the USDA-ARS SCINet Ceres high-performance computing cluster https://scinet.usda.gov. Each method was given a single 2.4GHz core with 16GB of RAM per replicate.

### 2.9 Performance comparison with Treemmer strain selections

An existing gene selection approach was introduced by Menardo *et al*., 2018, and we compared the runtime of *parnas* with this method. Treemmer is a randomized method without an explicit objective function. Therefore, we evaluated the representatives computed by Treemmer based on how close, on average, a taxon was to its closest representative (i.e., the k-medoids objective). For a simulated tree with 1,000 leaves, we used (i) Treemmer and (ii) random sampling to select *k* = 10, 50, 250 taxa and evaluated the selected taxa using the k-medoids objective. As Treemmer is randomized, for each *k* we performed random and Treemmer sampling 100 times to obtain a representative distribution.

### 2.10 Influenza A virus in swine dataset collection and curation

Influenza A viruses (IAV) are the causative agents of an important viral respiratory disease in pigs and humans. In pigs, subtypes of H1N1, H1N2, and H3N2 are endemic in swine around the world. Despite only three circulating subtypes, the genes encoding the surface glycoproteins, hemagglutinin (HA), and neuraminidase (NA), exhibit significant diversity due to two-way transmission of IAV between swine and humans (Nelson *et al*., 2012). Globally, approximately 30 phylogenetic clades of H1 and H3 genes were detected worldwide in the past 3 years in swine (Anderson *et al*., 2020), and across the same time period 16 H1 and H3 clades were detected in the United States (Arendsee *et al*., 2021). We downloaded 4,090 H1 and 1,572 H3 IAV in swine hemagglutinin (HA) genes from the Influenza Research Database (Zhang *et al*., 2017) [accessed on 06/13/2022]. We restricted analyses to sequences within the USDA influenza A virus in swine surveillance system (indicated by a nine digit alpha-numeric ‘A0’ code in the strain name) collected between 2016 and 2021. Phylogenetic clade classifications were determined using the H1 swine influenza H1 HA clade tool on IRD (Anderson *et al*., 2016) and H3 clades were assigned using octoFLU (Chang *et al*., 2019). The sequences were subsequently aligned using MAFFT v7.475 (Katoh and Standley, 2013) and we inferred H1 and H3 phylogenetic trees using FastTree v2.1.11 (Price *et al*., 2010) that were then rooted using Tree-Time v.0.8.4 (Sagulenko *et al*., 2018). We extracted the HA1 subunit amino-acid sequences using flutile v0.13.1 (https://github.com/flu-crew/flutile) since the HA1 subunit is a major target for protective antibody immunity and divergence in HA1 may act as a proxy for the antigenic distance between HA genes (Zeller *et al*., 2021).

## 3 Results

### 3.1 PARNAS outperforms existing representative sampling methods

#### 3.1.1 Comparison with ADCL

We compared the scalability of *parnas* with two core algorithms from Matsen *et al*., 2013. These algorithms solve the unweighted version of Problem 1 (representative sampling). The first algorithm, *ADCL-full*, is an exact algorithm for Problem 1, while the second algorithm, *ADCL-PAM*, is a heuristic and an adaptation of the classic Partition Around Medoids (PAM) k-medoids heuristic (Kaufman and Rousseeuw, 1990). *parnas* solves a more general problem (Problem 2) than ADCL methods in polynomial-time. Specifically, we measured both computational and memory savings in terms of the reduction of the size of the dynamic programming (DP) table.

*parnas* was significantly more scalable than ADCL methods on simulated data (Fig. 1). This is particularly encouraging in case of ADCL-PAM, as it is an inexact heuristic, whereas *parnas* is guaranteed to compute optimal representatives. Notably, *parnas* showed nearly linear runtime increase under both fixed *n* (number of taxa) and *k* (number of representatives). We further evaluated the DP table reduction due to Observation 1 and Lemma 2 above. For fixed *k* = 100, the average memory savings were 99.62% across 80 replicates. In particular, Observation 1 reduced the DP table size by 97.84% on average and Lemma 2 further decreased the size of the remaining table by 80% on average

**Figure 1:**
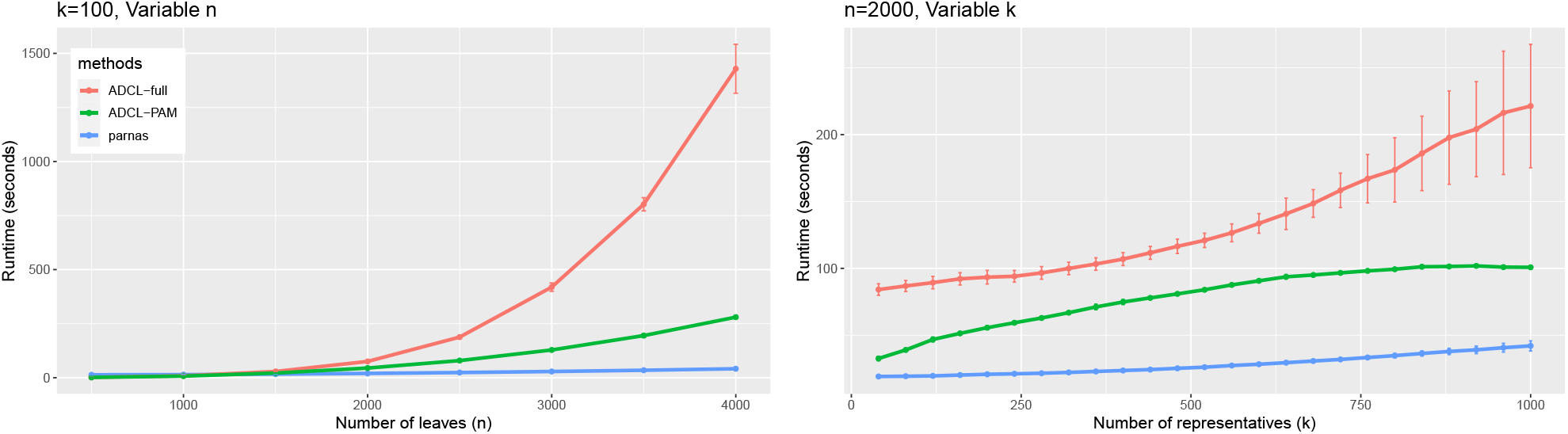
Runtime of *parnas* and ADCL methods with increasing number of leaves (left) and increasing number of representatives (right). The vertical error bars show standard deviation (across 10 replicates) around the mean.

#### 3.1.2 Comparison with Treemmer strain selections

We evaluated *parnas* against Treemmer in terms of the runtime and the representativity of generated selections. Treemmer uses a randomized procedure to select representative taxa across a phylogeny with a goal to reduce branch redundancy and does not explicitly solve the k-medoids problem like *parnas* and ADCL. We evaluated Treemmer in terms of the average distance of taxa to their closest Treemmer-selected representative (i.e., the k-medoids objective), and we computed how far Treemmer-selected representatives were from the optimal representatives by *parnas* in terms of that objective. *parnas* is significantly faster than Treemmer on large phylogenetic trees (Fig. 2). We observed that Treemmer representatives were generally better than randomly selected representatives, but they were 10-50% away from the optimum computed by *parnas*. That is, Treemmer-selected representatives are 10-50% further away from the taxa than the optimal representatives. The difference was particularly large for *k* = 250, where Treemmer representatives were 40-50% divergent from the optimum.

**Figure 2:**
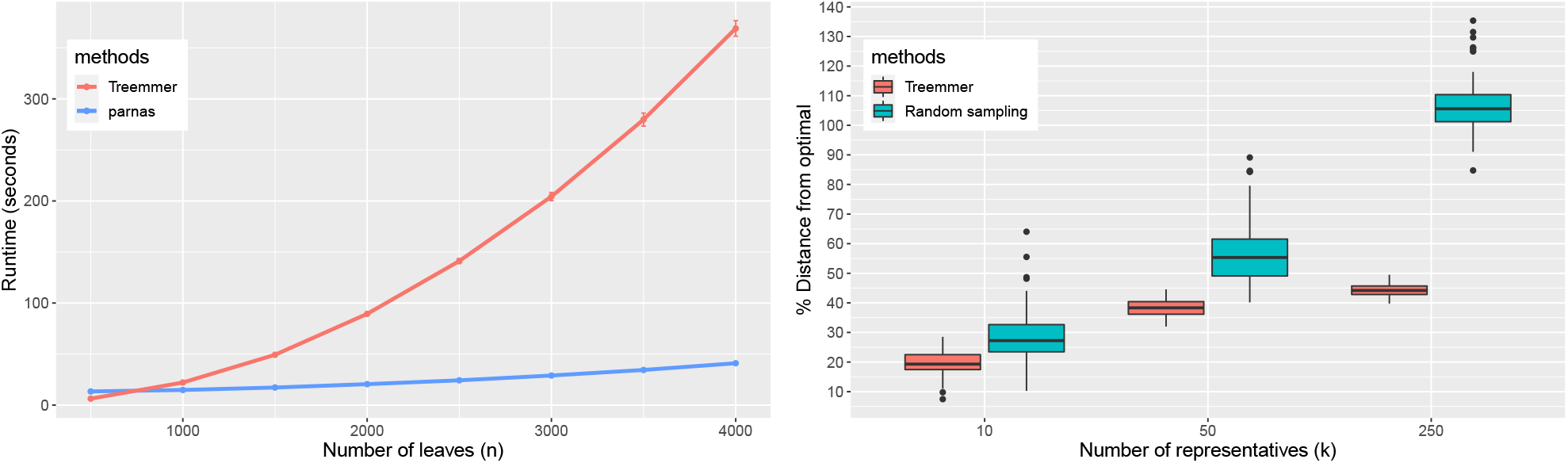
Runtime of *parnas* and Treemmer with increasing number of leaves and fixed *k* = 100 (left); and the assessment of Treemmer representatives in comparison to the optimal representatives computed by *parnas* and random sampling (right). supporting the effectiveness of our approach.

### 3.2 PARNAS-selected representatives in influenza A virus in swine

A central question in the development of a polyvalent influenza vaccine is determining how many HA components are sufficient to cover observed diversity (Anderson *et al*., 2012). We addressed this using the last 6 years of USDA IAV in swine surveillance data, and evaluated how many H1 and H3 HA genes were required to cover the genetic diversity of IAV in swine. Though the link between genetic diversity and antigenic phenotype varies (Bolton *et al*., 2019), we applied the conservative assumption that a hemagglutinin (HA) gene would retain some cross-reactivity against another HA gene that is within 5% amino-acid divergence in the HA1 subunit (Anderson *et al*., 2020). A second consideration in vaccine design, is the determination of when to update the components, and our study evaluated how long *parnas*-selected strains were adequate representatives of the more contemporary viruses detected in the USDA IAV in swine surveillance system.

For the H1 subtype, we used *parnas* to select 2, 4, and 6 of the most representative HA genes for the surveillance data collected in 2016, 2017, and 2018. Subsequently, we calculated how many HA genes in each year were within 5% divergence from the *parnas* representatives, i.e., for the 2016 selections, we determined how much diversity they covered in each year between 2016 to 2021. Similarly, for the H3 subtype, where fewer genetic clades cocirculate than in the H1 subtype (Arendsee *et al*., 2021), we used *parnas* to select 1, 2, and 3 of the most representative HA genes for 2016, 2017, and 2018. We executed *parnas* with the option to rescale the tree edges with the number of HA1 amino acid substitutions and specified a 5% divergence radius (16 amino acid substitutions).

We observed that the *parnas* selected representatives came from the most frequently detected circulating HA clades. For example, four 2017 selections from the H1 tree came from clades 1A.3.3.3 (*γ*), 1A.1.1 (*α*), 1B.2.2.1 (*δ*_1*a*_), 1B.2.1 (*δ*_2_), which were the most frequently detected H1 clades in the US that year (Arendsee *et al*., 2021). Similarly, selecting two representatives in the H3 tree consistently produced strains from the 1990.4.a and 2010.1 clades – the two most frequently detected H3 clades in the US since 2015 (Zeller *et al*., 2018).

Selecting four H1 HA genes were sufficient to cover over 70% overall diversity for the first year and the subsequent year, i.e., selections were adequate to cover the majority of observed diversity across two years (Fig. 3). Increasing the number of representative strains to six, guaranteed over 85% coverage in the first two years; whereas decreasing the number of selections to two reduced coverage to less than 50%. For the H3 subtype, three HA genes were sufficient to cover more than 85% of diversity in the same year, and the coverage remained consistently high for each subsequent year (Fig. 3). The 2016 representatives provided close to 90% coverage until 2020 (i.e., for 5 years straight). In H1s, the decrease in coverage was more pronounced over the years.

**Figure 3:**
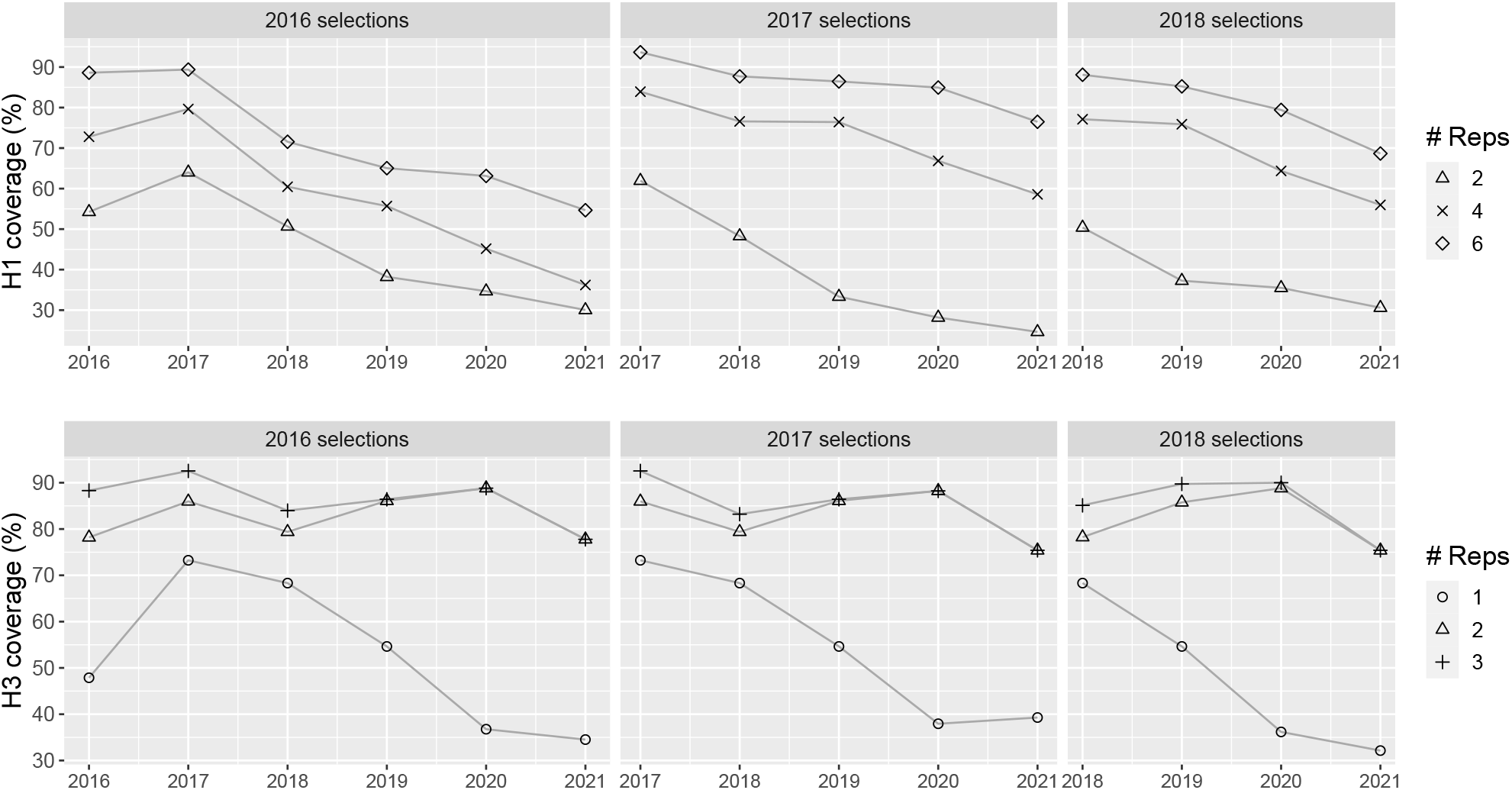
Percent of circulating influenza A virus in swine hemagglutinin genes that are within 5% divergence of the *parnas*-selected representatives. For H1 (top row), we selected 2, 4, and 6 representative strains with *parnas* for 2016, 2017, and 2018, and tracked how these representative HA genes covered H1 HA genes that circulated in the following years. Similarly, for H3 (bottom row), we selected 1, 2, and 3 representatives, and tracked how these representative HA genes covered H3 HA genes that circulated in the following years.

## 4 Discussion

Given the rapid growth of sequence data in genetic databases, representative subsampling techniques are essential for computation-intensive bioinformatics studies, objective selection of pathogen strains for phenotypic characterization, and for genomic epidemiology (Hill *et al*., 2021). There has been a surge in the number of tools that can parse sequence data or phylogenetic trees. One group of methods identify clusters of related sequences, e.g., TreeCluster (Balaban *et al*., 2019) or PhyCLIP (Han *et al*., 2019), but these are unable to objectively select strains within the identified clusters. A second group of methods select or remove single strains: such as TARDiS (Marini *et al*., 2021) that can perform time-aware sampling of genetic sequences, or Treemmer (Menardo *et al*., 2018) that reduces taxa on a phylogeny through pruning redundant branches. These selection approaches, either do not account for evolution, or are not able to objectively select representatives across tens of thousands of taxa in a reasonable time. *parnas* allows researchers to identify diversity groups within their data and objectively choose representatives. Further, *parnas* provides wide flexibility in constraining the pool of potential representatives through the specification of prior representatives or through the use of an optional coverage radius. Consequently, the method can achieve time-representative and geo-representative sampling by individually sampling from different time-periods and geographic regions across thousands of taxa.

We demonstrated that *parnas* is faster and broader in scope than ADCL (Matsen *et al*., 2013). Apart from subsampling and reference strain selection applications, ADCL was suggested to be used for genotype imputation (Kang *et al*., 2015; Ye *et al*., 2019). Therefore, *parnas* can be applied in a similar manner in the genotype imputation pipelines with large reference datasets. Matsen et al. also noted that “the computational complexity class of the ADCL optimization problem is not yet clear”. In this work we resolve this question and demonstrate that *parnas* solves the ADCL optimization problem, and a significantly more general problem, in polynomial time.

A primary motivation for developing *parnas* was the objective and representative selection of IAV in swine for phenotypic characterization. To this end, we demonstrated that *parnas*, unsupervised, selects representative strains from the most frequently detected IAV sequences in swine HA clades. We showed that as few as 6 HA genes (4 H1 and 2 H3) may be sufficient for effective representation of circulating IAV in the US for approximately two years. A consistent challenge in vaccine design is to determine what antigenic components are required, and *parnas* provides an objective approach to select HA genes that represent circulating diversity. Similarly, the algorithm provides a metric to determine when genetic coverage is reduced. Alternative pipelines for selection of representative strains in IAV research have previously involved (i) clustering of genes/genomes under a fixed divergence threshold and (ii) selecting consensus or random strains within the clusters (cf. Jones *et al*., 2021). The advantage of *parnas* is that it automatically achieves the same goal as these manual approaches, while also using an objective criterion that accounts for the interaction between the selected strains.

An additional application of *parnas* is the identification of genes that are not within a prescribed radius of prior representatives. For IAV in swine, the utility is the automated identification of HA genes that are genetically, and potentially antigenically novel. There are as many as 30 genetic and antigenically distinct clades of IAV in swine globally (Anderson *et al*., 2020), and more than 1000 IAV in swine HA genes are sequenced every year in the United States (Arendsee *et al*., 2021). *parnas* provides a rapid, reproducible, and objective approach to determine which of these viruses should be characterized using *in vivo* and *in vitro* methods. Linking computational assessments of circulating viruses with antigenic characterization can provide empirical data for use in pandemic risk assessments of viruses circulating in animal hosts (e.g., Souza *et al*., 2022). Further, *parnas* provides a convenient way to screen representative viruses for mutations in antigenic epitopes (Koelle *et al*., 2006; Bush *et al*., 1999; Plotkin *et al*., 2002) relative to existing viruses and vaccines as these changes may be associated with antibody-binding and IAV fitness (Łuksza and Lässig, 2014). This tool provides a rational and reproducible approach for parsing genomic surveillance data and developing a prioritization of strains to be comprehensively evaluated with a goal to detect novel viruses that may impact animal and human health.

## Acknowledgments

We gratefully acknowledge pork producers, swine veterinarians, and laboratories for participating in the USDA Influenza A Virus in Swine Surveillance System and publicly sharing sequences in NCBI GenBank.

